# TEsmall identifies small RNAs associated with targeted inhibitor resistance in melanoma

**DOI:** 10.1101/359471

**Authors:** Kathryn O’Neill, Wen-Wei Liao, Ami Patel, Molly Gale Hammell

## Abstract

MicroRNAs (miRNAs) are small 21-22nt RNAs that act to regulate the expression of mRNA target genes through direct binding to mRNA targets. While miRNAs typically dominate small RNA transcriptomes, many other classes are present including tRNAs, snoRNAs, snRNAs, Y-RNAs, piRNAs, and siRNAs. Interactions between processing machinery and targeting networks of these various small RNA classes remains unclear, largely because these small RNAs are typically analyzed separately. Here we present TEsmall, a tool that allows for the simultaneous processing and analysis of small RNAs from each annotated class in a single integrated workflow. The pipeline begins with raw fastq reads and proceeds all the way to producing count tables formatted for differential expression. Several interactive charts are also produced to look at overall distributions in length and annotation classes. We next applied the TEsmall pipeline to small RNA libraries generated from melanoma cells responding to targeted inhibitors of the MAPK pathway. Targeted oncogene inhibitors have emerged as way to tailor cancer therapies to the particular mutations present in a given tumor. While these targeted strategies are typically effective for short intervals, the emergence of resistance is extremely common, limiting the effectiveness of single-agent therapeutics and driving the need for a better understanding of resistance mechanisms. Using TEsmall, we identified several microRNAs and other small RNA classes that are enriched in inhibitor resistant melanoma cells in multiple melanoma cell lines and may be able to serve as markers of resistant populations more generally.

## Introduction

microRNAs (miRNAs) are small 21-22 nucleotide RNA molecules which have been shown to play a critical role in metazoan development and gene regulation, and while canonically derived from short hairpin RNA precursors, have been shown to originate from a variety of sources including tRNAs and introns.^1,2^ In addition to governing development, sRNAs play a critical role in repressing transcripts derived from repetitive regions of the genome like transposons. In animals, siRNAs and piRNAS function to repress transposons in somatic cells, and the germline respectively^3,4^. Identification of miRNAs and siRNAs which originate from non-canonical regions of the genome is more challenging with few programs designed to detect sRNAs from all classes in both unique and repetitive genomic loci. It is for this reason we present TEsmall, a package designed specifically for the simultaneous analysis of sRNAs derived from a variety of genomic features. In particular, this package facilitates the discovery of intriguing biological phenomena otherwise masked by insufficient annotation of repetitive genomic elements, such as siRNAs, and allows these elements to be easily incorporated into downstream differential analysis through packages like DESeq2^5^.

We have tested the ability of TEsmall to characterize the expression profiles of small RNAs from a variety of classes in the context of melanoma cell lines responding to targeted inhibitors of the BRAF oncogene. The genetic basis of melanoma development is fairly well understood, with activating mutations in the oncogene BRAF occurring in a majority of melanoma patient tumors^6^, which also harbor hundreds of secondary mutations of unknown impact. Specific inhibitors that target activated BRAF as well as the downstream MAPK/ERK signaling pathway have been developed, which dramatically reduce the growth of melanoma cells in patients. However, the effects of these drugs typically extend patient lifespan for six months or less, as the tumors rapidly develop resistance to these targeted therapies^7^. While some tumors resistant to BRAF inhibitors acquire additional genetic lesions that elevate MAPK or AKT signaling^8^, many therapy-resistant cell lines establish resistance without a clearly understood mechanism of resistance^9^. Changes to small RNA profiles in melanoma cells responding to targeted inhibitors is an especially poorly understood subset of the genomic and transcriptomic changes that occur. To understand how small RNA alterations might contribute to the development of resistance to BRAF inhibitors in 451Lu melanoma cells that carry BRAFV600E mutations, we undertook a small RNA sequencing study of cells before and after the establishment of BRAF inhibitor resistance.

## Materials and Methods

### Melanoma cell culture

In this dataset, 451 Lu patient derived melanoma cell lines were used to explore the sRNA profiles of cells that are either sensitive or resistant to small molecule inhibitors of the BRAF kinase. Specifically, the melanoma patient derived 451Lu-Par cells are grown in standard growth media (DMEM with 10% FBS), while the 451Lu-BR cells are grown in standard growth medium supplemented with a 1uM concentration of the BRAF inhibitor vemurafenib. Both cell lines are adherent cells grown in standard 2D cell culture. The derivation of BRAF inhibitor resistance in these cells lines is described by Villanueva et al.^7^ and the cell lines are available from Rockland for both 451Lu cells (cat: 451Lu-01-0001) and 451Lu-BR cells (cat: 451Lu BR-01-0001).

### Small RNA sequencing libraries

Total RNA was extracted using the Ambion PureLink RNA Mini Kit to extract up to 2 µg of total RNA from ~1×10^6^ melanoma cells from either the 451Lu-Par or 451Lu-BR lines. Following Bioanalyzer verification of RIN numbers at or above ~9, the RNA extracts were next used to create small RNA sequencing libraries. The small RNA sequencing libraries were prepared with the Illumina TruSeq Small RNA Library Preparation Kits using an input of 1.2 µg total RNA and following the manufacturer’s protocols as described, using 15 PCR cycles to reduce the likelihood of PCR amplification artifacts. The libraries were pooled and indexed with 6nt Illumina barcodes, such that 6 libraries could be sequenced per lane on an Illumina Genome Analyzer IIx. The reads were sequenced as single-end 50bp reads, to a depth of approximately 35 million reads per library. The dataset is available through GEO at the following accession number: GSE116134. A table of sequenced and mapped read counts for each library is presented in Supplemental File 1.

### qPCR Validation

Taqman qPCR assays were used to validate the analysis results of TEsmall for a subset of microRNAs. Specifically, standard Taqman qPCR probes were obtained for the following microRNAs: miR-100, miR-184, and miR-211. Control probes were obtained for RNU58 and the U6 small RNA. Custom Taqman probes were obtained for the predicted mature sequence of the novel candidate miRtron derived from the VIM intron 6 locus. The Taqman protocol was followed exactly as described, using 1 µg of total RNA as the input from two biological replicates of the 451Lu-Par and 451Lu-BR cell lines, and using 3 technical replicates per biological replicate. The “Comparative Ct” analysis method described in the manufacturer’s protocol was used for calculating fold change, standard deviation, and t-test based P-values. Briefly, the three technical replicates for each probe were combined to create a mean Ct value per probe per sample. The average of the Ct values from the two control probes in each sample was then subtracted from each microRNA Ct value to create a normalized “∆-Ct” value for each microRNA in each sample. Following averaging of ∆-Ct values between the two biological replicates in each condition, the ∆-∆-Ct value was calculated as the difference in mean ∆-Ct values for the same microRNA across conditions. Fold change represents 2^∆-∆-Ct^, with errors on each Ct value combined quadratically.

### TEsmall module

TEsmall functions by accepting raw input in FASTQ file format from next generation sequencing platforms in conjunction with genomic annotation sets via an online server. Adapters associated with siRNA library preparation are trimmed by TEsmall through the cutadapt package^10^. In order to remove degradation products from abundant ribosomal RNAs, rRNA derived reads are next filtered from the data before proceeding to analysis. This mapping step allows for up to 2 mismatches and filters a single alignment per read specified by the option: bowtie -v 2 -k 1 using bowtie (v1.2.1)^11^. Small RNA reads remaining after rRNA filtering are then aligned more stringently, disallowing mismatches, option: bowtie -v 0 -a -m 100. All alignments in this step which map to fewer than 100 genomic loci are reported allowing for the classification of multimapper reads common to sRNA data, in particular structural RNAs like tRNAs and transposable element targeting siRNAs. Following alignment to the genome, each alignment annotated via a sequential decision tree, as follows. The reads are aligned to each annotation category in order, then removed from the pool of alignments in order to facilitate priority annotation of, for example, intronic microRNA reads that should properly be annotated as microRNAs rather than intronic RNAs. The default order is: structural RNAs, miRNAs and hairpins, exons, sense transposons, antisense transposons, introns, and ultimately piRNAs. This ordering, can be re-ordered by the user to suit the application and user preferences about annotation priority. An HTML output file is then created using python based Bokeh tools^12^ to visualize the abundance distributions, length distributions, and mapping logs of all small RNAs in the dataset (Fig. 1). In conjunction with this HTML output, TEsmall compiles multiple flat text output files, including a counts file that is structured to be directly compatible with DESeq2^5^ for differential analysis. The abundance calculations for these counts files are 1/n normalized, where n represents the number of alignments per read, to ensure no double-counting of multimappers.

**Figure 1.**
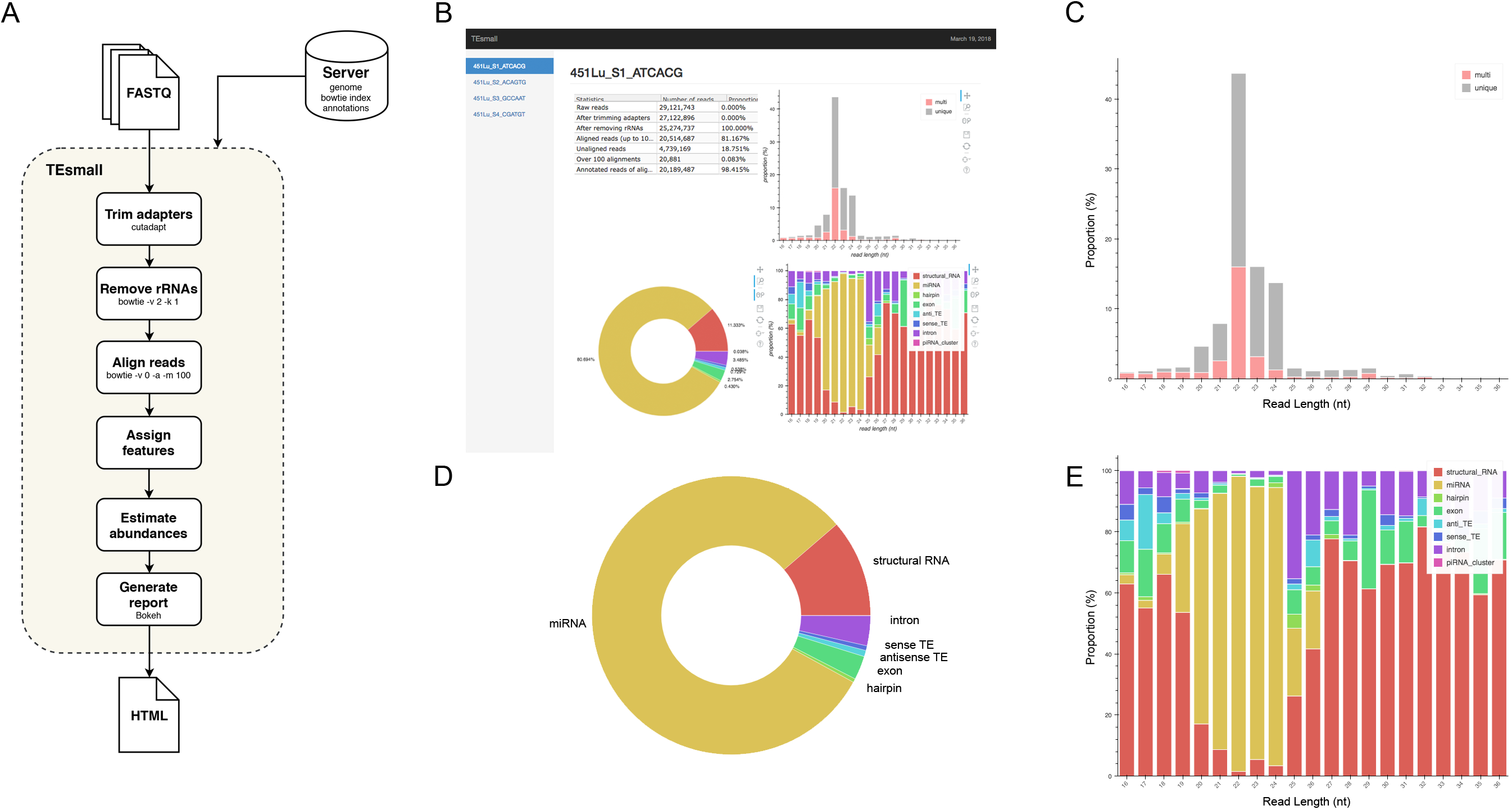
Flow chart and output HTML of TEsmall. (A) Flow chart of TEsmall’s treatment of input high-throughput sequencing data, input genome indices, and output. (B) Screenshot of HTML output file for one sample (C) Bar plot depicting size distribution of unique and multimapper reads. (D) Circle plot depicting distribution of reads to each subtype. (E) Bar plot depicting proportion of subtypes across read length.

The TEsmall code is available open-source from GitHub at the following location:https://github.com/mhammell-laboratory/tesmall.

### Differential Analysis with DESeq2

The counts file produced by TEsmall were subsequently imported into DESeq2 (v1.18.1) to perform differential analysis between 451Lu-PAR and 451Lu-BR cell lines, as follows. The counts file was filtered to remove low abundance species (< 20 counts across all libraries) and increase the sensitivity of DESeq2. Normalization of the counts for differential analysis was performed using the default DESeq2 method during statistical analysis. For downstream visualization, the counts were normalized by the built-in variance stabilizing transformation (VST) method in DESeq2. Small RNAs with an adjusted P-value < 0.05 were considered statistically significant. The full DESeq2 output file is given in Supplemental File 2.

### Visualization

Figures were produced using the R packages ggplot2^13^, gplots^14^ and GenomicRanges^15^ for scatterplots, heatmaps, and wiggleplots respectively. Python package matplotlib^16^ was used for all barplots. RNA secondary structures were rendered using the forna webtool^17^, secondary structure for the Arg-ACG-1-2 tRNA was pulled from the UCSC GtRNAdb tRNA covariance model, and structure of vimentin intron 6 was predicted using RNAfold’s minimum free energy model.^18^

## Results

### TEsmall Workflow

TEsmall is a package specifically designed to identify sRNAs derived from a variety of genomic features simultaneously, such that users can evaluate the relative abundances and profiles of many source of small RNAs on a common scale in a single pipeline. This serves as a novel improvement to currently available software such as mirDeep2^19^ and piPipes^20^, which are optimized for the analysis of miRNAs and piRNAs respectively, but are not equipped to evaluate both types of small RNAs together. TEsmall is also designed so that its output is optimally formatted for downstream differential analysis with statistical modeling software, such as DEseq2^21^. A flowchart describing the entire TEsmall workflow is given in Figure 1A, with example output charts given in panels 1B-E. Specifically, in the first module of TEsmall, raw small RNA sequencing reads from Illumina NGS sequencing platforms serve as the input without the need to pre-process the data before beginning analysis. TEsmall first trims adapter contaminants from the reads and then filters the reads for appropriate size ranges, with a default of 16-36 nucleotides in length. The next module of TEsmall removes contaminating ribosomal RNA fragments by mapping with bowtie^11^ to a library of rRNAs for the specified genome. Removal of rRNA reads is critical as rRNA degradation products are a major source of contamination for sRNA data. Remaining reads are mapped to the genome, with a default mapping strategy optimized for repetitive regions with up to 100 alignments per read, though this may be altered by the user. The reads are next sequentially annotated to several small RNA classes and genomic features, with a decision tree implemented to prioritize annotation categories. This has the goal of attributing reads mapping to intronic microRNAs as “microRNA” reads, for example, rather than annotating these reads as having an intronic source. Following annotation of each read, aggregate abundances are calculated for each sequencing library and output as a counts table suitable for downstream differential expression analysis. Importantly, any multimapper reads in these counts tables are weighted according to the number of genomic loci from which they derive (1/n where n is the number of alignments) to avoid any double-counting of multimapper reads in the counts tables. In addition to an output file including all raw count data per sample, RNA species ID, and type classification, TEsmall provides an aesthetic output HTML (Fig. 1B) summarizing distribution of read lengths (Fig. 1C), proportion of reads assigned to each sRNA type (Fig. 1D), and distribution of reads of a particular size to each of the sRNA categories (Fig. 1E). In addition to these summary plots, TEsmall presents a table with summary statistics of read proportion, raw input and trimmed read counts to quickly assess any potential biases in library preparation that may affect downstream normalization.

### Application of TEsmall to melanoma sRNA profiles

As described, drug resistance is a known hurdle in the treatment of melanoma, driving the need for a better understanding of how cells develop resistance. We have chosen to investigate the alterations in sRNA profiles, as one marker of cellular state. To investigate the effect of BRAF kinase drug resistance on sRNA composition in patient derived melanoma cell lines, we performed differential expression analysis following classification by TEsmall. Resistant lines were derived through exposure of 451Lu patient derived parental cell lines to increasing concentrations of vemurafenib up to 1uM. Resistant clones were selected and expanded before exposure to an increase in vemurafenib. Cells were otherwise treated as described in Villanueva et al.^7^ Raw count data was normalized as described in Materials and Methods by DESeq2. All VST normalized counts were averaged between parental or resistant replicates and plotted against each other to visualize trends of expression across sRNA subtypes without filtering for significant or abundant transcripts, significantly differentially expressed transcripts are represented by solid coloring (Fig. 2). Overall, there appears to be a trend towards lower expression of many sRNA classes in the 451Lu BRAF resistant samples (Fig. 2 and Fig. 3B). Upon filtering for the most abundant (base mean across all replicates > 500) and significantly differentially expressed transcripts (multiple-hypothesis testing adjusted p-value < .05), trends of lower sRNA expression in the BRAF resistant samples was still seen for many intronic, exonic, and transposon mapped sRNA species (Fig. 3). Interestingly, miRNAs show an even distribution of species with negative and positive log fold changes, and since miRNAs were the most abundantly sequenced small RNAs in the libraries, this rules out a normalization issue as the explanation for down-regulated small RNAs in the other classes. It may be of interest that, after filtering for significance and abundance, structural RNAs with a negative log fold change are exclusively tRNAs with the exception of vaultRNA 2-1, and those with a positive log fold change are almost exclusively snoRNAs. Details of the particular sRNAs differentially expressed in each of these classes are given in Supplemental File 2.

**Figure 2.**
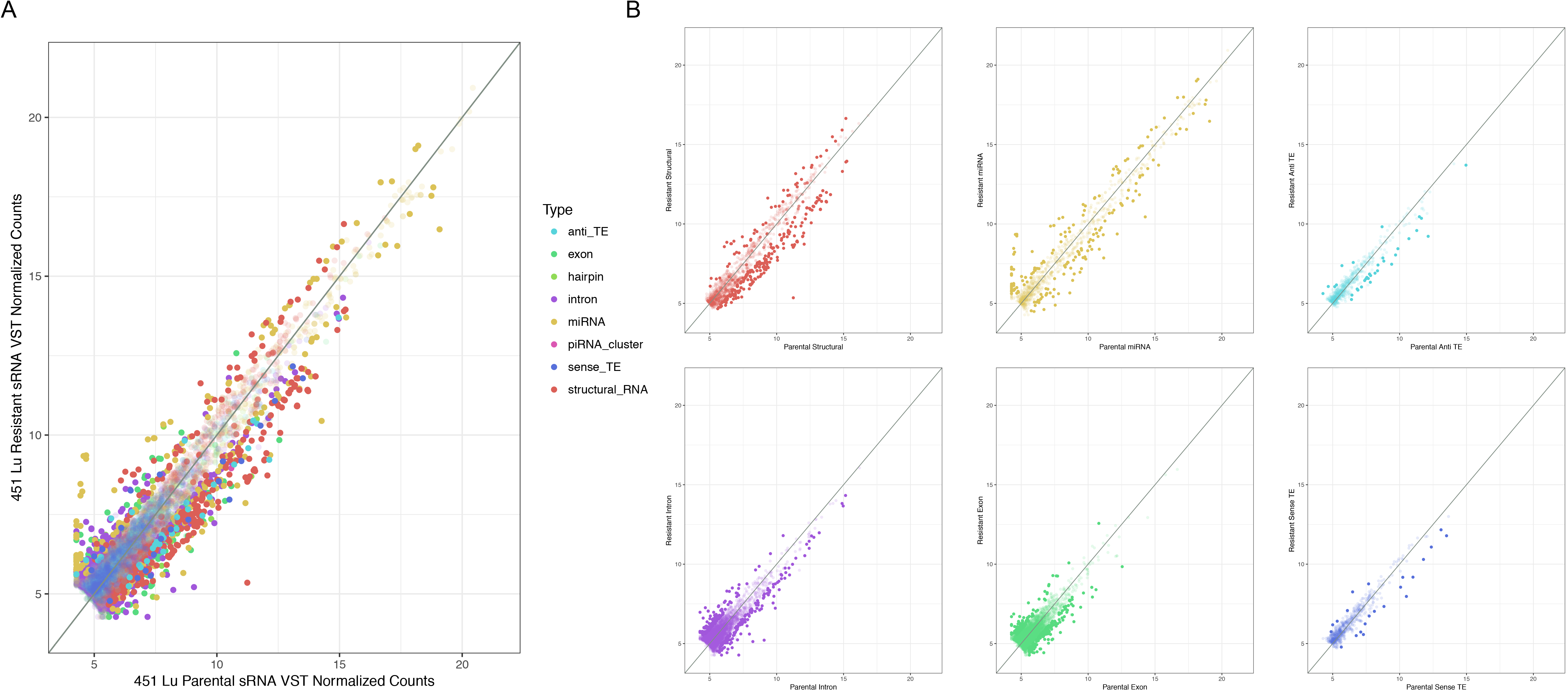
Scatterplots depicting BRAF resistant versus parental RNA VST normalized counts. (A) Overlay of all subtype scatterplots. (B) RNA subtype specific scatterplots. Transparent points represent RNA species with adjusted p-value <. 05.

**Figure 3.**
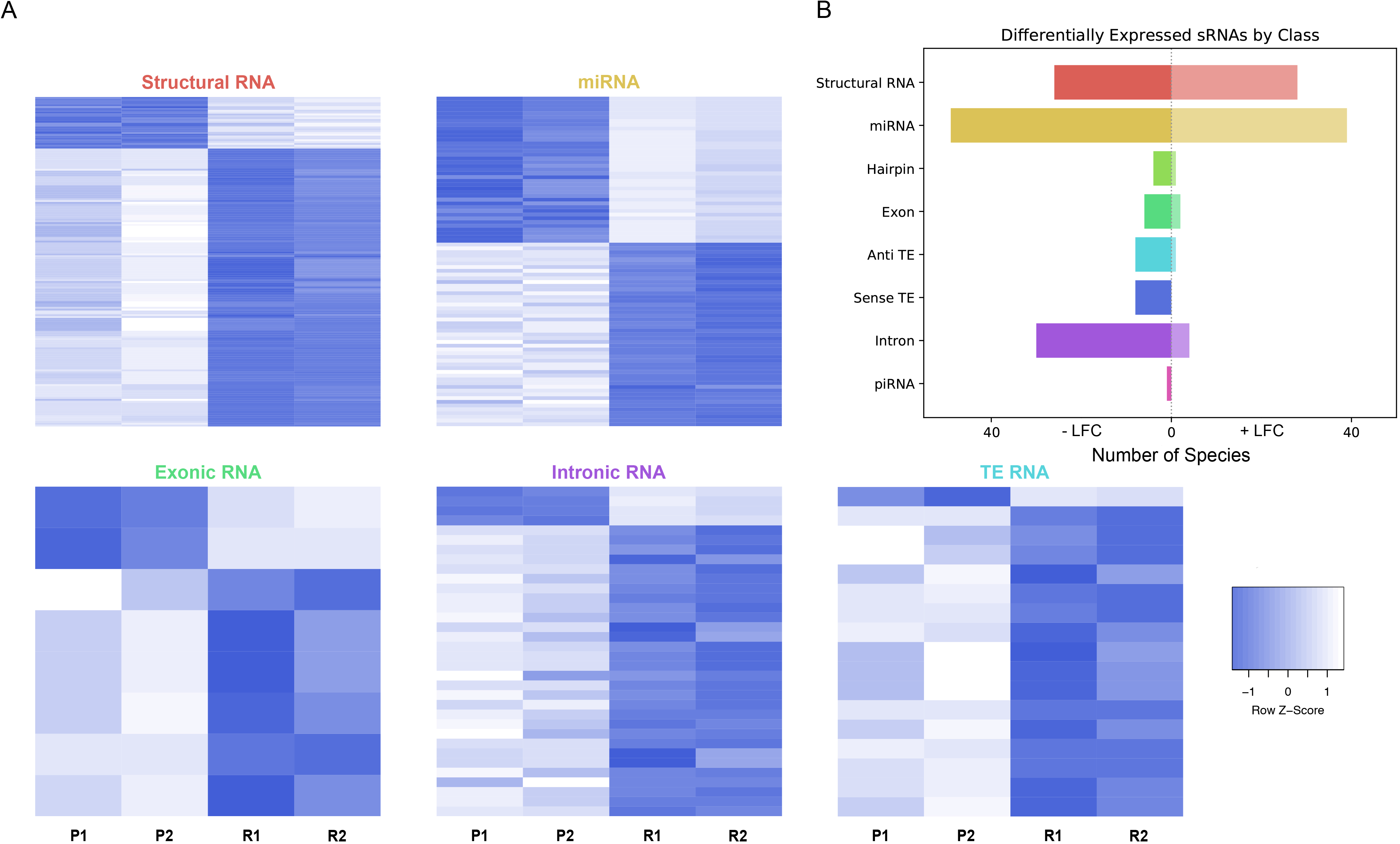
Differential expression of highly abundant transcripts. (A) Heatmaps depicting all significantly differentially expressed genes per subtype, all heatmaps are normalized to between -1 and 1 by row. (B) Bar plot depicting number of abundant and significantly differentially expressed species with a negative or positive log fold change per subtype. Structural RNA species are collapsed on duplicate tRNAs. Abundance is determined by a base mean between samples of greater than 500 counts. P1, P2, R1, and R2 represent parental replicates 1 and 2 and BRAF resistant replicates 1 and 2 respectively.

### Type specific analysis of sRNA species and Validation with qPCR

Several miRNAs which are significantly differentially expressed in our dataset have been previously described in the literature as playing critical roles in melanoma progression or epidermal differentiation. This includes the miRNAs miR-184, miR-211, and miR-100. In other contexts, miR-184 has been shown to arrest epidermal differentiation through derepression of Notch in normal human keratinocytes and murine epidermis^22^. While expression of Notch in keratinocytes is known to have a tumor suppressive phenotype, its expression has the opposite effect in melanocytes through upregulation of the PI3K/Akt and MAPK pathways^23^. Our data showed an approximate 5-fold increase in miR-184 expression in BRAF inhibitor resistant cells in comparison to parental (Fig. 4A and Supplemental File 2), consistent with a model where MAPK pathway activation provides a mechanism for BRAF inhibitor resistance.^7,24^ It has also been shown that BRAF inhibitor resistance can be mediated by regulatory escape of the transcription factor MITF from the MAPK pathway, where MITF overexpression itself conferred resistance in several melanoma cell lines.^25^ Consistent with a high MITF state, our data shows a significant upregulation of miR-211, derived from the MITF activated gene melastatin, and a significant downregulation of miR-222, known to be inversely correlated with MITF expression.^26^ Finally, miR-100 was also shown to be significantly downregulated in our data; this was a miRNA of interest as it has been implicated in prostate cancer as a repressor of the oncogene mTOR.^27^

**Figure 4.**
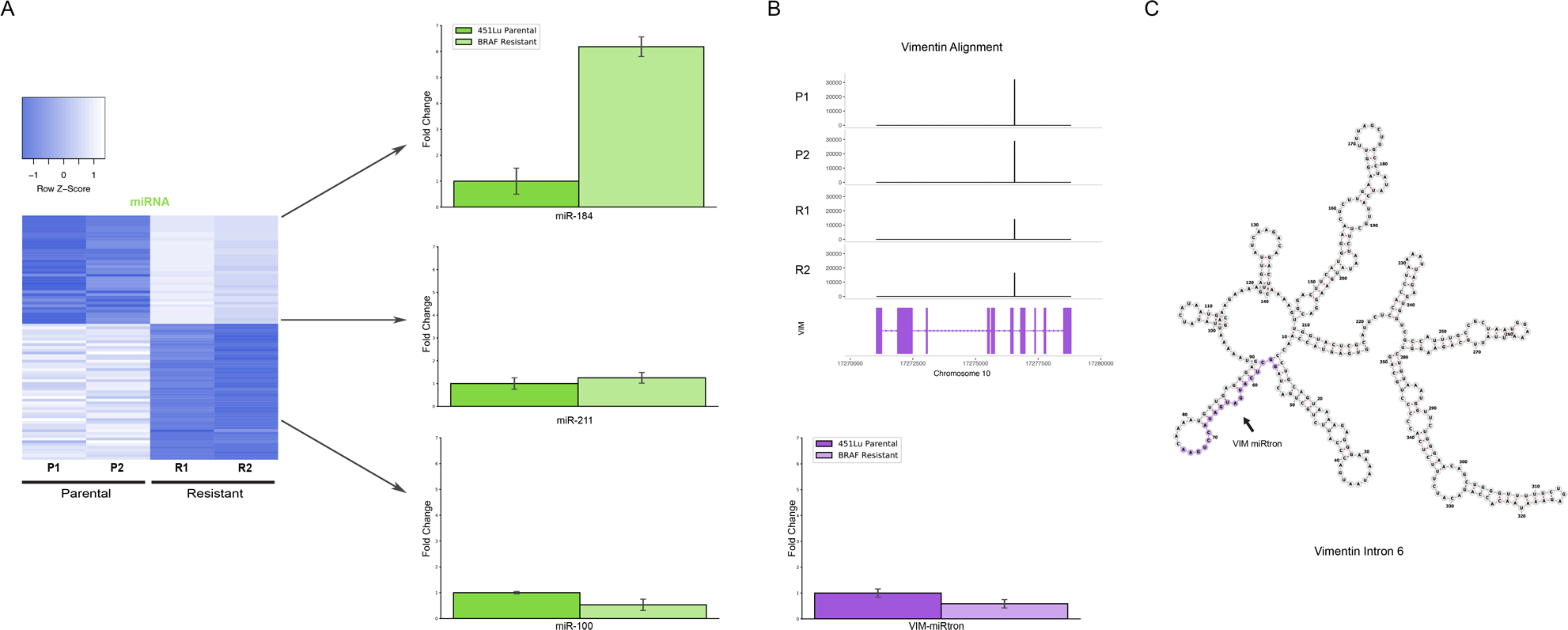
Detailed analysis miRNAs of interest. (A) qPCR representing log fold change of miRNAs 184, 211, and 100 in respect to parental expression levels across 3 replicates. (B) qPCR representing log fold change with respect to parental cell line and BAM gene alignment tracks across samples of the VIM miRtron C) RNAfold predicted secondary structure of VIM intron 6 with miRtron highlighted in purple.

To validate the expression profiles from our TEsmall based differential expression analysis, we performed qPCR on several miRNAs of interest including miRNAs miR-184, miR-211, and miR-100, all of which recapitulated the trend observed in our small RNA-seq dataset (Fig. 4A). Reassuringly, the expression alterations of miR-204 and miR-211 seen in our data were also seen in an alternative melanoma derived cell line A375 in Díaz-Martínez et al. following induction of BRAF inhibitor resistance.^28^

Upon further investigation into individual RNA species from different subtypes for follow up, we encountered an interesting and novel 21 nucleotide sRNA associated with the sixth intron in the vimentin gene (VIM) (Fig. 4B). Intronic microRNAs that derive from short spliced introns with internal hairpin structures have been observed previously and dubbed miRtrons.^29^ MiRtrons are typically generated via the splicing machinery and subsequently processed by DICER in the cytoplasm, bypassing the canonical nuclear Drosha processing steps. As visible in the minimum free energy secondary structure prediction by RNAfold, the candidate miRtron of interest is located in a stem loop structure which appears conducive to processing by DICER (Fig. 4C). The length of this mature sRNA, its abundance as a single RNA species, and its secondary structure within VIM intron 6 are all consistent with miRtronic miRNAs. This is particularly interesting as VIM is a known marker for the epithelial to mesenchymal transition and is well expressed in many cell types, but has not previously been shown to harbor a miRtron, suggesting this VIM miRtron might represent a novel miRNA with particular abundance in melanoma cells.

In addition to miRNAs, TEsmall recognizes several other types of sRNAs. It has been previously reported that tRNA derived small RNA molecules (tRFs) can silence LTR retrotransposable elements through occupation of the primer binding site (PBS) as an adaptation of the role of tRNAs as retroviral primers.^2,30^ Through TEsmall, one is able to detect reads associated with tRNAs and transposable elements in the same pipeline facilitating observation of phenomena such as these. In our analysis, several species of sRNAs mapping antisense to transposable elements were significantly depleted in BRAF resistant cell lines compared to parental (Fig. 2). Upon further investigation we were able to determine that a subset of these reads were tiRNAs derived from the Arg-CGY family of tRNAs (Fig. 5B). These candidate tiRNAs mapped to a subset of HERVs including HERV3, HERV30, MER51, and others (please see Supplemental File 3). This is consistent with previous literature showing HERV-R retrotransposons are primed by Arg tRNAs (Fig. 5A) and tRNA derived fragments (tRFs) can occupy retroviral primer binding sites to suppress transposon activity.^2,30^ It is important to note that the Arg-CGY sRNAs reported by TEsmall are consistent with the tRFs previously described in Schorn et al. as they are 18nt CCA-appended fragments originating from the 3’ T-arm of tRNAs.^2^ This is shown graphically in Fig. 5, where the pileup of reads at an example HERV PBS locus can be seen in Fig. 5B, and the pileup of these same reads at the originating tRNA locus can be seen Fig. 5C-5D. In the tRNA profiles, other tRNA fragments can be seen outside of the tiRNA generating 3’ end, but these do not predominantly accumulate as a single abundant sRNA species.

**Figure 5.**
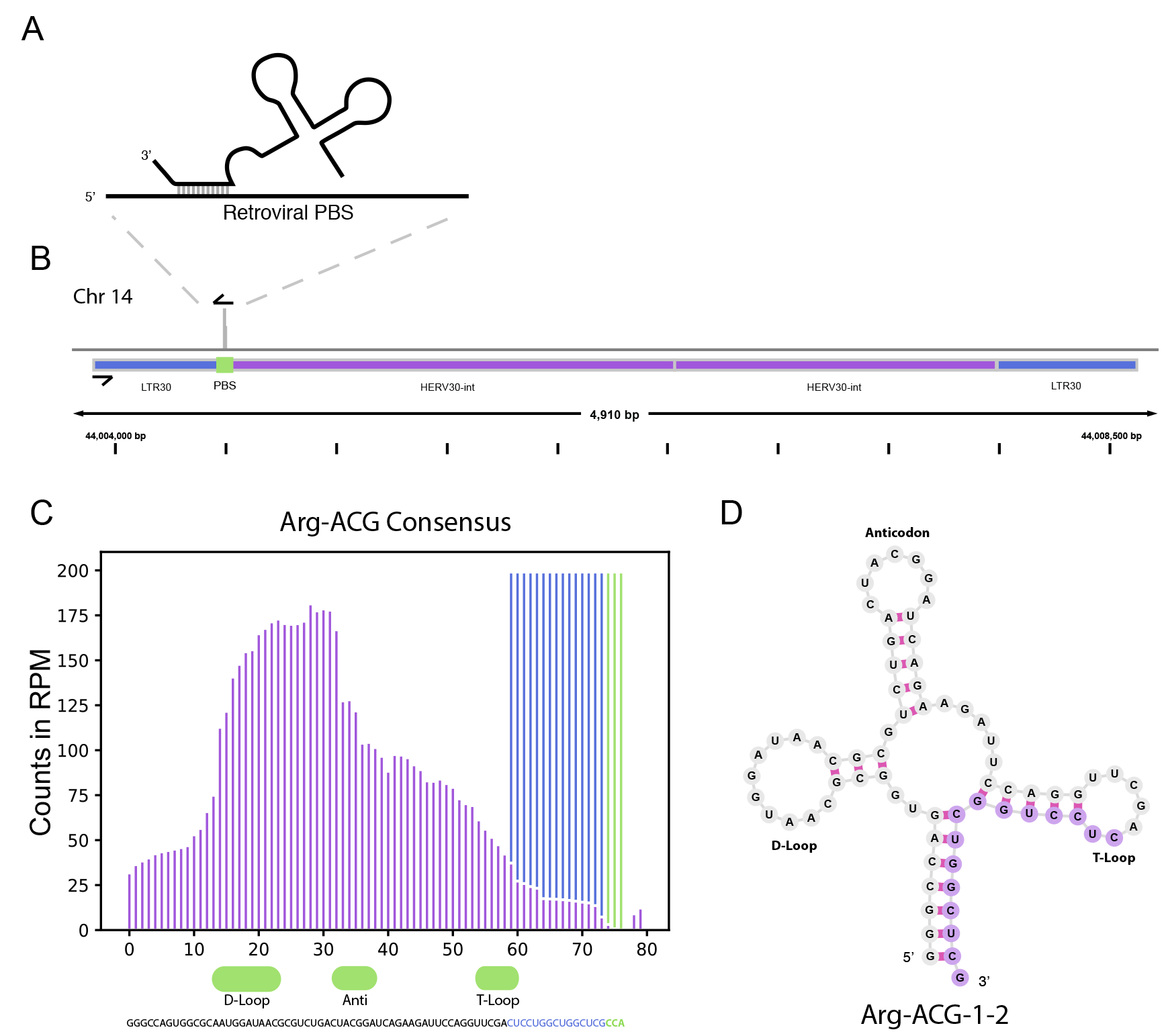
tRNA and TE interaction through primer binding sites. (A) Diagram of primer binding by tRNA to facilitate retroviral reverse transcription. (B) Read alignment track of Arg-CGY family derived 18 nt CCA tailed fragment to HERV30 PBS. (C) Consensus histogram of reads distributed from Arg-CGY tRNAs, and derived tiRNAs D) Secondary structure of Arg-CGY family member Arg-ACG-1-2 with highlighted 15 nt CCA (-) fragment.

In addition to the miRNAs and tiRNAs highlighted above, several additional species of small RNAs were reported by TEsmall as differentially expressed in the 451Lu BR cells including: siRNAs mapping to transposable element loci, exonic loci, and a variety of structural RNA classes. The structural RNA group included snoRNAs, snRNAs, tRNA fragments, and a vault RNA. The full list of differentially expressed small RNAs can be found in Supplemental File 2.

### Comparison of TEsmall with sRNA Analysis Software

Several software packages exist to characterize small RNA data for expression profiling analysis. However, programs designed for this purpose such as miRDeep2^19^ and ShortStack^31^ typically focus on a particular category of sRNA, predominantly miRNAs. One package that considers multiple sRNA types is piPipes^20^, a software suite designed for both piRNA and siRNA analysis. While piPipes functions well to annotate and characterize piRNAs by read pileups associated with the ping-pong cycle of piRNAs, it is not particularly suited for annotation of sRNAs from other types of genomic loci, such as miRNAs, siRNAs, and tiRNAs. PiPipes will provide plots of read distribution across lists of transposable elements and piRNA clusters, however, one cannot access tables of these counts with associated TE annotation, suitable for differential expression analysis. While piRNAs are annotated with their respective piRNA clusters, siRNAs are assigned a chromosomal coordinate providing some difficulty in determining patterns in the sources or targets of these reads. It is also of import that intron derived miRNAs like the VIM miRtron were not captured, as there is no mechanism by which to assign siRNA reads beyond mapping the chromosomal coordinates associated to preloaded annotation sets associated with TEs and piRNA clusters. TEsmall, which does not perform piRNA-specific ping-pong analysis, provides a complementary package that is designed to be a general purpose small RNA analysis suite that can identify and analyze many types of sRNAs concurrently, presenting the output in a format intended for expression profiling analysis.

## Discussion

TEsmall is a software package with novel functionality in that it allows the user to simultaneously map and annotate many types of sRNAs including structural RNAs, miRNAs, siRNAs, and piRNAs. This allows one to compare trends in expression between all sRNA types and investigate the cross-talk between distinct sRNA regulatory pathways. Other packages released to date focus on individual sRNA types like miRNAs^19,31^ or piRNAs^20^ and while optimized for these applications, are not adapted for comparison across sRNA categories. In addition to handling multiple classes of sRNAs, the output of TEsmall is formatted for direct integration into downstream analysis pipelines. TEsmall’s output files are compatible with statistical analysis software like DESeq2 and efficient heatmap generation. In addition to requiring little data preprocessing, TEsmall outputs an aesthetic HTML file of charts (Fig. 1B) which allows for fast and effortless assessment of library quality, sRNA composition, and size distribution. TEsmall can also be expanded to function for any novel sRNA species provided the appropriate annotation files are available, allowing it to serve as a powerful tool to study RNA biology in many organisms.

We applied TEsmall to a novel dataset in which we compared the effects of BRAF inhibitor resistance on sRNA abundance in melanoma derived cell lines. In this analysis we found several microRNAs whose expression was altered in BRAF inhibitor resistant cells in comparison to parental lines. A table of these hits can be found in Supplemental File 2. Among these candidates, we experimentally validated changes in expression of miRNAs miR-184, -211, and -100. Of particular interest is the novel Vimentin derived miRtron candidate, whose expression pattern was also experimentally validated. Close examination of the characteristic read pile up associated with the VIM miRtron, and secondary structure of intron 6 are all consistent with miRtron processing pathways. Further investigation will be required to determine if this is a true miRtron formed through an intermediate spliceosome derived lariat independent of the Drosha microprocessor subunit, or is instead a canonical Drosha-dependent miRNA.

In addition to revealing miRNAs previously described in the literature, TEsmall detected several novel classes of small RNAs which would not have been found using packages designed for miRNA analysis. TEsmall allows the user to investigate tRNA derived fragments which have been shown to play a critical role in LTR retro-transposon suppression^2^. In the melanoma dataset, we identified a novel candidate tiRNA that appears to derive from ARG-tRNAs and to potentially regulate several HERV-R type LTR elements through occupancy of the primer binding site. Other types of siRNAs that regulate transposon expression were also shown to be differentially expressed in these datasets, suggesting the possibility that transposon-derived transcripts are altered in these BRAF inhibitor resistant melanoma cells.

It is well known small noncoding RNAs of different subtypes types work in conjunction to regulate cellular processes through complex networks, particularly in the realm of transposon silencing. piRNAs known to regulate transposon expression in the germline have been found to work in cooperation with siRNAs to perform this task.^32^ In plants, miRNAs have been shown to play a role in transposon silencing by serving as an intermediate to form 21 nucleotide siRNAs via RNA dependent RNA polymerase and while the mechanism would be disparate from plants, hints of miRNAs facilitating transposon silencing have been seen in animals as endogenous and introduced retroviral elements with homologous regions to miRNAs have lower genomic activity.^33,34^ Current sRNA analysis packages are specific to one or two types of sRNAs making it easy to overlook biologically interesting patterns of interaction between sRNA classes. For this reason, we have created TEsmall, an easy to use package with aesthetic output designed for the concurrent expression analysis of multiple sRNA subtypes.

## Acknowledgements

MGH is a scholar of the Rita Allen Foundation. KO acknowledges funding from the NIH training grant 2T32GM065094-16. We would also like to acknowledge the Meenhard Herlyn lab for help, advice, and for the kind gift of the 451Lu cells.

## Author Contributions Statement

Authors KO and MGH analyzed the data, generated the figures and wrote the manuscript. MGH and AP designed the experiments and generated the data. WWL wrote the code for the TEsmall pipeline, packaged the code for GitHub, and contributed to the writing of the manuscript.

